# ViRNN: A Deep Learning Model for Viral Host Prediction

**DOI:** 10.1101/2024.03.30.587436

**Authors:** Pierre Sphabmixay, Blake Lash

## Abstract

Viral outbreaks are on the rise in the world, with the current outbreak of COVID-19 being among one of the worst thus far. Many of these outbreaks were the result of zoonotic transfer between species, and thus understanding and predicting the host of a virus is very important. With the rise of sequencing technologies it is becoming increasingly easy to sequence the full genomes of viruses, databases of publicly available viral genomes are widely available. We utilize a convolutional and recurrent neural network architecture (ViRNN) to predict the hosts for the *Coronaviridae* family (Coronaviruses) amongst the eleven most common hosts of this family. Our architecture performed with an overall accuracy of 90.55% on our test dataset, with a micro-average AUC-PR of 0.97. Performance was variable per host. ViRNN outperformed previously published methods like k-nearest neighbors and support vector machines, as well as previously published deep learning based methods. Saliency maps based on integrated gradients revealed a number of proteins in the viral genome that may be important interactions determining viral infection in hosts. Overall, this method provides an adaptable classifier capable of predicting host species from viral genomic sequence with high accuracy.

## 1. Introduction

With the rise of numerous viral outbreaks in the past ten years, including the recent SARS-CoV-2 pandemic, understanding the determinants of virus-host interaction will be key for predicting and understanding the spread of such viruses. There are now rich databases of information to explore for clues to viral pathogenesis by looking through viral genome sequences. Viral genomes encode numerous proteins important for the lifecycle of the virus, including capsid and envelope proteins for structure and cell entry, as well as various DNA or RNA interacting proteins for genome replication. The functionality of these proteins can help determine the viral life cycle, including which hosts the virus can infect. For example, the spike protein (S) of coronaviruses like SARS-CoV-2 is a critical protein for host infection and interspecies transmissions as it codes for the surface protein that mediates both cell attachment and membrane fusion. Spike protein variations can lead to changing cell tropism, allowing for alternative hosts. This protein is perhaps the most well studied in terms of host cell tropism, however, it is possible that other aspects of the viral genome sequence could hint towards potential hosts for a virus.

Various supervised and unsupervised learning techniques have been employed in this field, including KNN based models, support vector machines, and recurrent neural networks, although no deep learning techniques have been applied to the coronavirus family ^1–3^, which has a particular affinity for zoonotic transfer between species, as demonstrated by recent outbreaks of SARS, MERS, and COVID19. Here we demonstrate the application of a combined convolutional and recurrent neural network architecture which identifies the actual host of a virus from genomic sequences between 11 host species with high accuracy.

## 2. Methods

### 2.1. Coronaviridae dataset

The coronavirus family consists of a number of viruses^4^ infecting a wide range of vertebrate hosts from bats to humans. They have been studied and sequenced for quite some time, leading to over 30,000 sequences being publicly available. For this project we curated data deposited in the NCBI GenBank under the taxon *Coronaviridae* (taxon ID: 11118). This project utilized a freeze created on 4/08/2020, containing all 33,618 sequences annotated with the txid 11118. Sequences were directly downloaded from the NCBI using publicly available webtools, packaged as part of the Biopython toolkit ^5^. Although over 33,000 sequences were available, not all of these were annotated with host sequences. After pruning 26,066 sequences (77.54%) of sequences remained, which contained sequences annotated with 797 unique hosts. As predicting this number of hosts would be untenable, and many listings were only semantically different (e.g. *Homo sapiens vs Homo sapiens-female)* we analyzed the most frequent hosts and found the 30 most abundant species listings comprised 78% of the data, with diminishing returns thereafter. We then combined functionally identical host names (e.g. combined pig, swine, and porcine into Pig) for ease of prediction to yield a total of 11 hosts for further analysis.

### 2.2. Data-preprocessing

As can be seen in Figure S1, there is a large discrepancy in the length of sequences for this family, with the larger sequences representing full genomes, and the smaller sequences representing only partially sequenced genomes or proteins. In order to standardize the input to our CNN+RNN architecture, which requires equal length sequences, we split each sequence into overlapping subsequences of length *k* with overlap *n*, various combinations of *k* and *n* were tested to optimize the model. Sequences shorter than *k* were zero-padded on the right. After generating subsequences, each sequence and resulting host was one-hot encoded and a TensorFlow Dataset was generated from this data. Training, testing, and validation sets were created using the *test_train_split* function in sklearn, with 64% of the data used to train, 16% to validate, and 20% to test. Class weights were calculated based on abundance to mitigate the class imbalance in the dataset using an sklearn function.

### 2.3. KNN and SVM Benchmarks

In order to compare our deep learning model to more common machine learning (ML) algorithms we implemented two other models based on previously published work that aimed at predicting hosts from viral genomic sequences : a K-Nearest-Neighbor (KNN) algorithm and a Support Vector Machine (SVM) algorithm for multi-classification.

KNN algorithms were established ^2^ based on unaligned genomic sequences for various viruses with various degrees of success. In order to perform KNN, distance matrices need to be computed to predict the nearest neighbors for any given point. To compute pairwise distance this methodology first requires the computation of k-mer profiles for each sequence ^6^. For a given k value, the number of words containing k letters is computed for each sequence and then stored as a profile using the python module kPal^6^. Because DNA/RNA contains 4 possible nucleotides, the number of features (words) for kProfiling increases rapidly with 4^k^. Various distance metrics between kProfiles have been established specifically for the purpose of genomic analysis. Common metrics such as Euclidean distance (**Eu**) and Manhattan distance (**Ma**) have been used in this context ^2^, showing better performance with the **Ma** distance. Other metrics have been developed and established: the Dissimilarity statistic (**D2**) ^7^ and the improved sheep statistic **D2*s** ^8^.

In order to produce the KNN model we then implemented this workflow:

- Computation of kProfiles for each sequences in our data (k=3,4,5,6,7 tested)
- Computation of distance matrices (n x n sequences at a time, n = 100).
- Establishment of KNN algorithm: measuring accuracy using a Leave One Out Classification Method.

An important observation is that this method was very computational expensive and increased dramatically with the number of sequences (See Supplementary Table 1). Ideally the distance matrices would be computed on the whole dataset for KNN but it would be practically not possible given the size of the dataset(>26,000 sequences). For this work we computed 100x100 matrices at a time. The maximal value for Li and Sun (2018) ^2^ was 707x707 for their study on Coronavirus.

**Table 1:** Per-Host Accuracy by sequence length

| Host | k=3000bp | Full Sequence | Number of Sequences (Test Set) |
| --- | --- | --- | --- |
| Bat | 85.81% | 91.73% | 336 |
| Camel | 74.6% | 77.86% | 128 |
| Cat | 84.09% | 96.80% | 216 |
| Chicken | 97.84% | 99.30% | 1277 |
| Cow | 72.2% | 88.88% | 96 |
| Dog | 61.74% | 88.54% | 99 |
| Duck | 71.19% | 80.48% | 53 |
| Human | 88.20% | 89.28% | 735 |
| Pig | 98.67% | 99.27% | 1065 |
| Pigeon | 66.67% | 82.85% | 43 |
| Yak | 84.38% | 91.66% | 24 |
| **Weighted Average** | **90.55%** | **95.3%** |  |

Another commonly used ML method to predict viral hosts from unaligned sequences is SVM on kProfiles ^3^. Because kProfiles have output dimensions too big for SVM (16384 features for 7-mers while the dataset only contains 26,000 sequences), a prior step of feature dimensional reduction is necessary. Principal component analysis is a useful method to reduce dimension while conserving variance from the dataset. For this work we implemented a C-classification task on a SVM on PCA components of the dataset using a radial base function as kernel for the SVM (See Figure S5 and S6 for PCA and SVM hyperparameter optimization).

### 2.4. Neural Network Architecture and Training

Our work was conducted with Python 3.5 and models were trained and evaluated using TensorFlow 2.1. Precision recall curves were generated using code derived from the sci-kit learn toolbox. Some code was derived from published code VIDHOP^1^ and examples given in the TensorFlow and Keras documentation.

The model utilized in this project was derived from a similar CNN+RNN architecture recently published (Figure 1)^1^. Briefly, we utilized two different models, with the only difference being the type of recurrent layer utilized, one used long-short term memory (LSTM) while the other used gated recurrent units (GRU).

**Figure 1:**
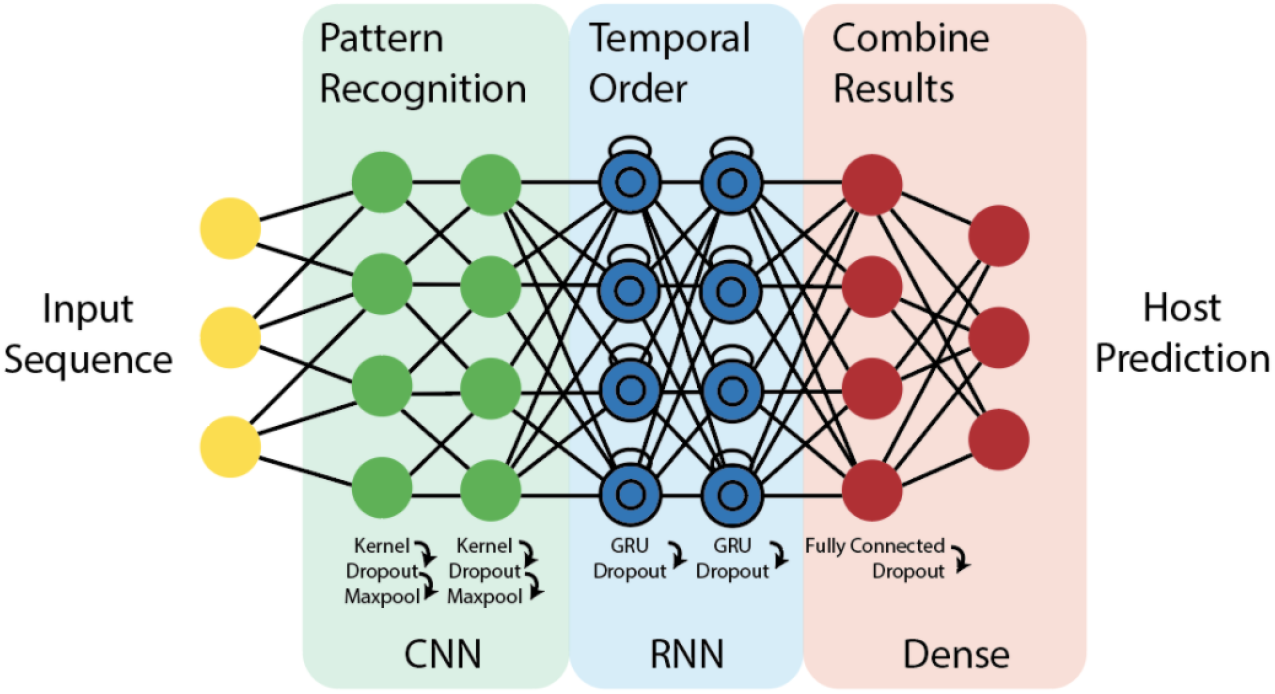
CNN+RNN Architecture. We utilized a CNN+RNN architecture derived from previous work^1^. We tested LSTM and Input GRU layers for the Sequence middle 2 RNN components of the model. Note this figure is derived from reference 1.

The model utilized two 2D-convolutional layers, with a kernel size of 1% of the input sequence length, stride 1, and a leaky ReLU activation. Each of these layers was followed by a max pooling layer with pooling size of 2 and stride 1. This fed directly into recurrent layers with sigmoid activation; either two Bidirectional LSTM layers (150 nodes) or two Bidirectional GRU layers (150 nodes), each followed by a dropout layer. 5% dropout was used to train all models. Following this there are two fully connected layers, the first with 150 nodes and ELU activation, followed by a dropout layer, and then the second with 11 nodes (number of hosts) and softmax activation. Categorical cross entropy was used as an objective function along with the Adam optimizer. Models were trained for 5-15 epochs depending on the purpose, with early stopping based on the validation loss to monitor for overfitting. Because of the dramatic class imbalance between many of the hosts (Figure S1, 43 times more sequences for the most abundant (chicken) compared to the least abundant (Yak)) we trained each model with class weights, calculated as the inverse of the fraction that host composed in the dataset.

### 2.5. Interpretability Methods

In order to understand how the model makes a prediction given an input sequence, we tried to implement various “block-box” interpretability methods for deep learning models. Various techniques such as DeepLIFT ^9^ have been successful at interpreting NN but are not straightforwardly compatible with RNN. For this work we implemented a saliency method known as Integrated Gradient in order to create a saliency map on each input sequence to probe for important regions/nucleotides that guided the model towards a specific host prediction.

Two different baselines were tested:

- N-nucleotide baseline where all the nucleotides are labeled as N (any nucleotide), the 5th channel in the 1-hot-encoding of the RNA sequences (A,C,T,G - N).
- An embedding of the average nucleotide distribution within the sequence.

Raw saliency maps produced by the Integrated Gradient method on a given sequence exhibited two peaks at both ends of the sequence (See Figures S8) which are thought to be caused by vanishing gradients in the RNN, a well known problem in the community.^10^. In order to circumvent this problem and create relevant saliency maps of the sequences we created a new workflow:

- Each sequence is truncated through a sliding window of fixed size (3000bp) that scans over the sequence (shift increment of 120bp)
- For each truncated sequence a saliency map is computed from the integrated gradient method
- Resulting aligned and shifted saliency maps are then average to produce an overall saliency map over the entire starting sequence.

Sequence logo maps were created with the package LogoMaker, and the software Geneious was used to visualize the viral genomes for protein annotation.

## 3. Results

### 3.1. Neural Network Optimization

Although many of the network parameters were borrowed from literature as previously described, we tested two different recurrent layer types (GRU and LSTM). We tested both architectures as GRU layers are reported to train faster and use fewer variables, simplifying the training. Only LSTM architecture has been used for this type of model in the literature. We also tested pre-processing our sequences into various *k* lengths, with a 10% overlap between all sequences. Each of these models was trained for 5 epochs using the Adam optimizer. We found a sharp decrease in the validation loss after 5 epochs at an input subsequence length of 3000-5000 bp (Figure 2). Shorter sequences did not perform as well, although the GRU model demonstrated lower loss after 5 epochs than the LSTM for all conditions tested. Note that no model was trained for sequence length 10,000 bp for the GRU model due to computational limitations. Although we observed slightly lower validation loss with the GRU model at 5000 bp length, we utilized the GRU model with input subsequence length 3000 bp for further training because it trained in a substantially shorter period of time.

**Figure 2:**
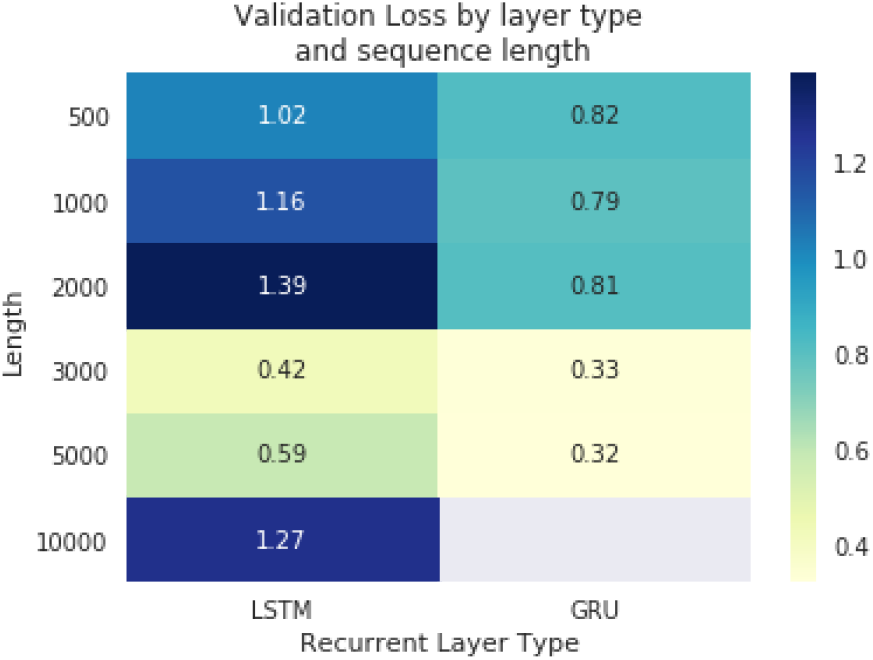
Validation loss after 5 epochs for models with either LSTM or GRU layers across various input sub-sequence lengths. No model was trained for GRU length=10000.

After identifying the optimal architecture, we trained the GRU based CNN+RNN model for 15 epochs with a batch size of 500 sequences. We utilized early stopping with a patience=3 epochs based on the validation loss to prevent overfitting.

We saw no improvement in loss after 7 epochs and thus training was ended and weights were restored to epoch 7. These data are visualized over 10 epochs in Figure 3. Note that the AUC is high even at the start of training due to the class imbalance.

**Figure 3:**
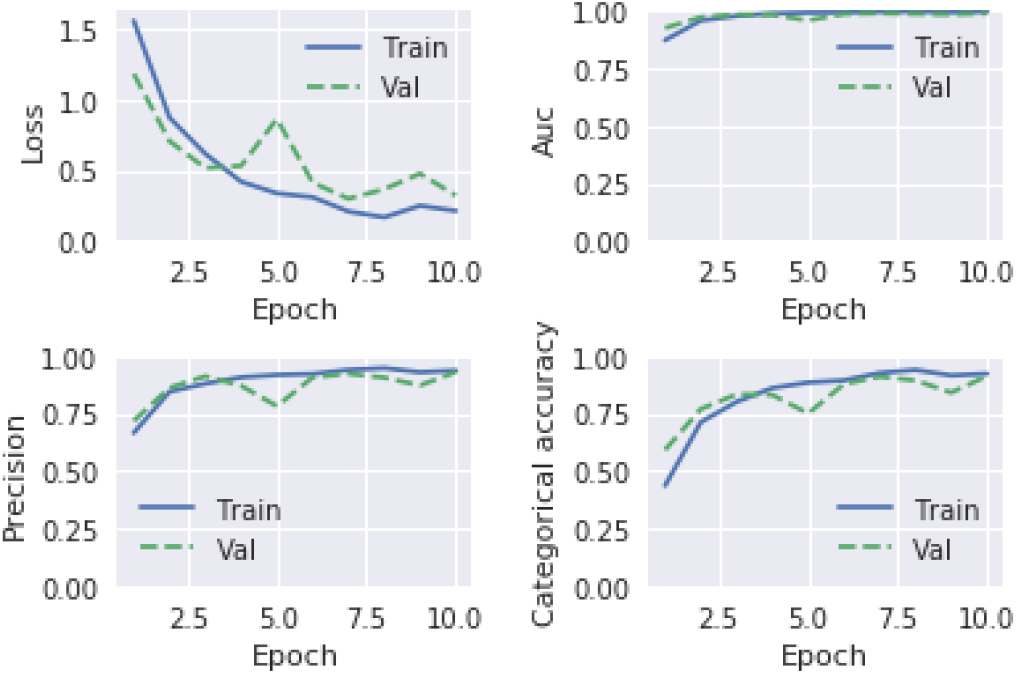
Training of CNN+RNN GRU based model. Training metrics plotted over 10 epochs for the training and validation sets, the loss function (upper left), the area under the curve (upper right), the precision (bottom left), and the categorical accuracy (bottom right).

### 3.2. Performance of RNN

Because of the dramatic class imbalance in our dataset, we chose to represent the performance of our model in precision recall curves to best demonstrate the accuracy of the model (Figure 4). Over the entire test dataset of 20360 sequences, the model performs with an average of 90.88% accuracy over all hosts when predicting from our test set consisting of subsequences of 3000 bp. Based on the precision-recall curve, the model performs well for hosts with a large number of sequences (Human, Pig, Chicken) and not as well for hosts with fewer sequences like dog. Notably, the hosts with the fewest sequences, Yak and Pigeon, had a higher categorical accuracy and AUC-PR compared to hosts with a higher number of samples like Dog and Duck. We find that the micro-average AUC-PR for the classifier is 0.97, with the most accurate host being for Pig (AUCPR=1) and the least accurate being for Duck (AUCPR=0.67). Figure 5 details the host predictions made for each actual host. For all hosts predicted the accuracy was greater than 60%.

**Figure 4:**
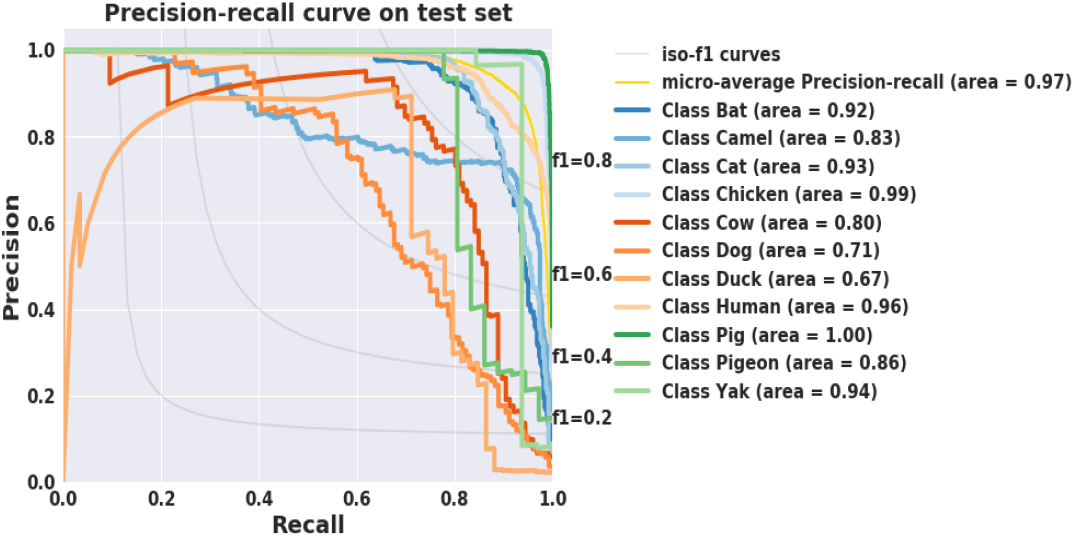
Precision-Recall Curve for testing dataset using ViRNN. Curves generated with sklearn’s precision recall function by class (e.g. class 1 versus rest) as well as a micro-average precision-recall curve for all classes. The AUC-PR for each class is listed in the legend.

**Figure 5:**
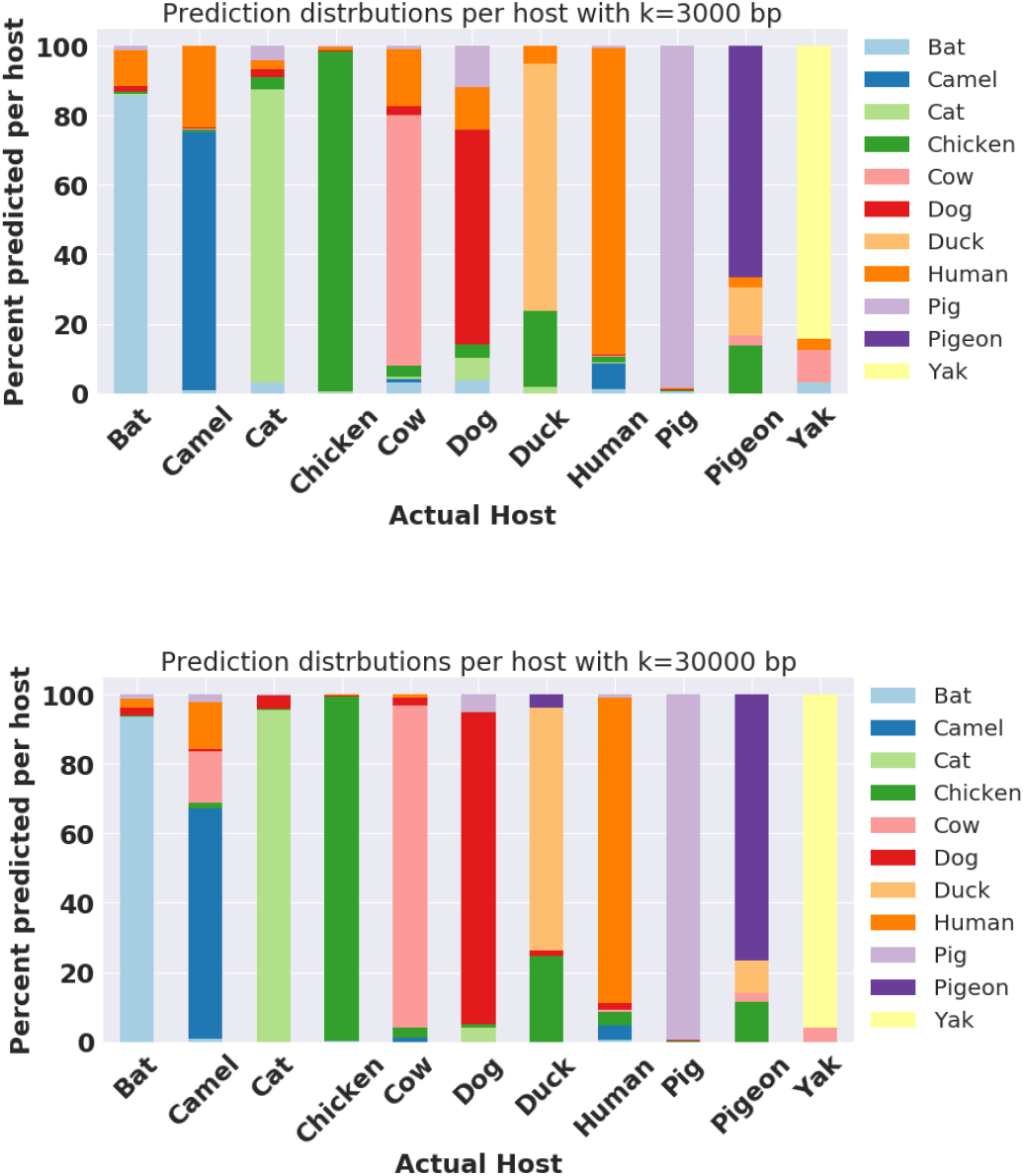
Model prediction by host. For each ground truth host the proportion of predictions for that host and all other hosts are indicated by the bar size. (TOP) for sequence length=3000 bp, (BOTTOM) for full sequence length from GenBank (no sub-sequencing)

We expected deviations from our sub-sequencing regimen to reduce the accuracy of the model since the model was not trained on these sequences. We were surprised to see the accuracy increase on the full sequence set (no sub-sequencing) to an average of 95.3% accuracy across all hosts. Since this dataset was very unbalanced, we looked more carefully at a host-by-host view of the predictions for more insight. We saw gains in accuracy across all 11 hosts (Table 1, Figure 5). Some gains were rather modest, with chicken increasing about 2% while others including dog and pigeon increased by almost 20% in accuracy with the full length sequences.

### 3.3. Generalizability of RNN on alternative viral taxon

Although our model was trained on the family of *Coronaviridae*, we were curious how generalizable the learned features were for viruses of other families. We chose to curate sequences from the *Orthomyxoviridae* family, a prevalent family of viruses including influenza which infects some of the same hosts as *Coronaviridae*. We downloaded 114,351 sequences from the NCBI GenBank taxon id: 11308, annotated with four hosts in common with our coronavirus dataset: pig, chicken, human, and duck. These were subsequenced with the same method as used to train our model (k=3000 bp) and the predictions were compared with the actual host information. We found the model performed poorly for the *Orthomyxoviridae* family, with an overall accuracy of 18.0 % on all of the sequences. Rather than predicting the actual host, it seems the model mostly generated predictions of the most abundant hosts including chicken, bat, and pig, although it seems for all hosts the model predicted the low abundance dog host for at least 10% of the sequences (Figure 6). Overall, these data indicate the model does not perform well on out-of-distribution virus families.

**Figure 6:**
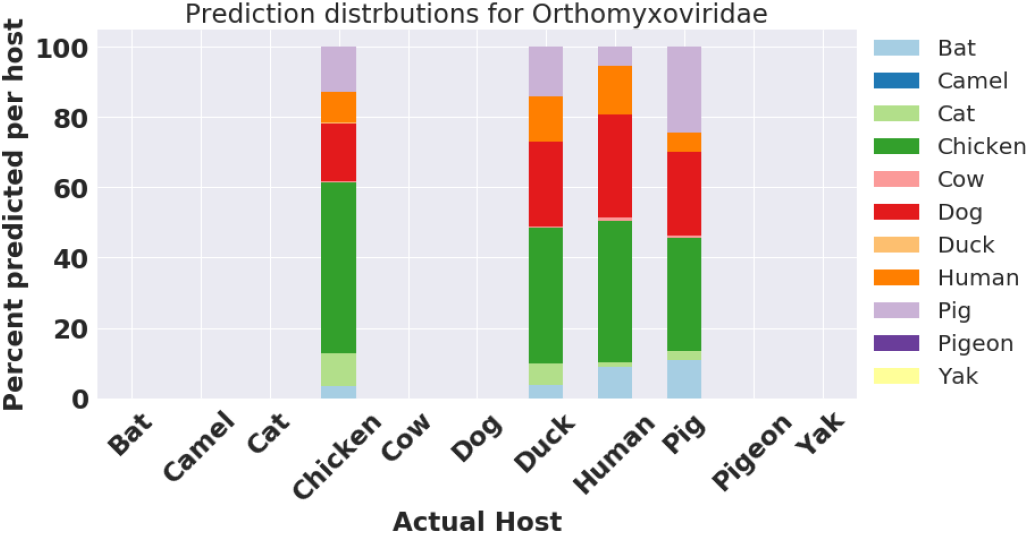
Model prediction by host for *Orthomyxoviridae*. For each host the proportion of predictions for that host and all other hosts are indicated by the bar size. Predictions conducted on subsequences of length k=3000bp.

### 3.4. Comparison of RNN with other methods

We created two other models with more canonical ML (KNN and SVM) methods that were previously published for the same objective: predicting viral host from an unaligned sequence of the virus RNA. Our goal is to compare performance of these two ML algorithms with our deep ViRNN.

First, we tested the prediction accuracy on the KNN model. Because this method was the most computationally expensive, we could not use all the dataset for the model (See Supplementary Table 1). With a restricted dataset the KNN achieved an accuracy of 0.74 ± 0.09 for the D2S* metric and using 1-neighboring threshold. (See Figure 7 left).

**Figure 7:**
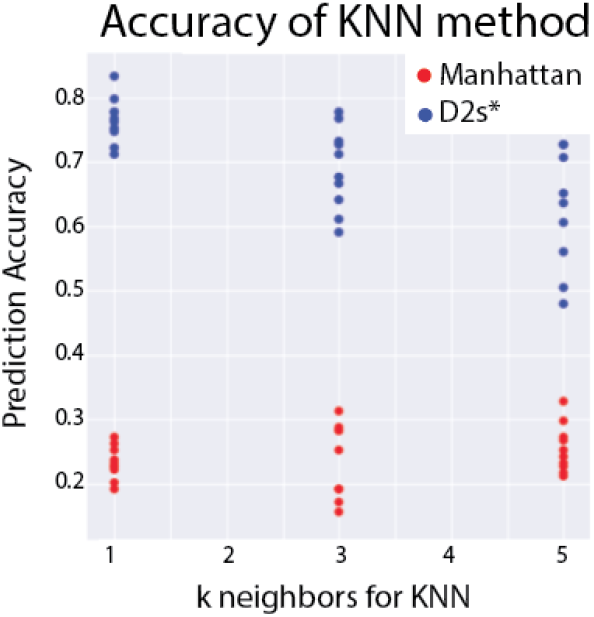
(left): Accuracy for KNN classification method. Two distance metrics (Ma and D2S*) Each point represents 100x100 samples picked at random (n=10 points)

**Figure 7.**
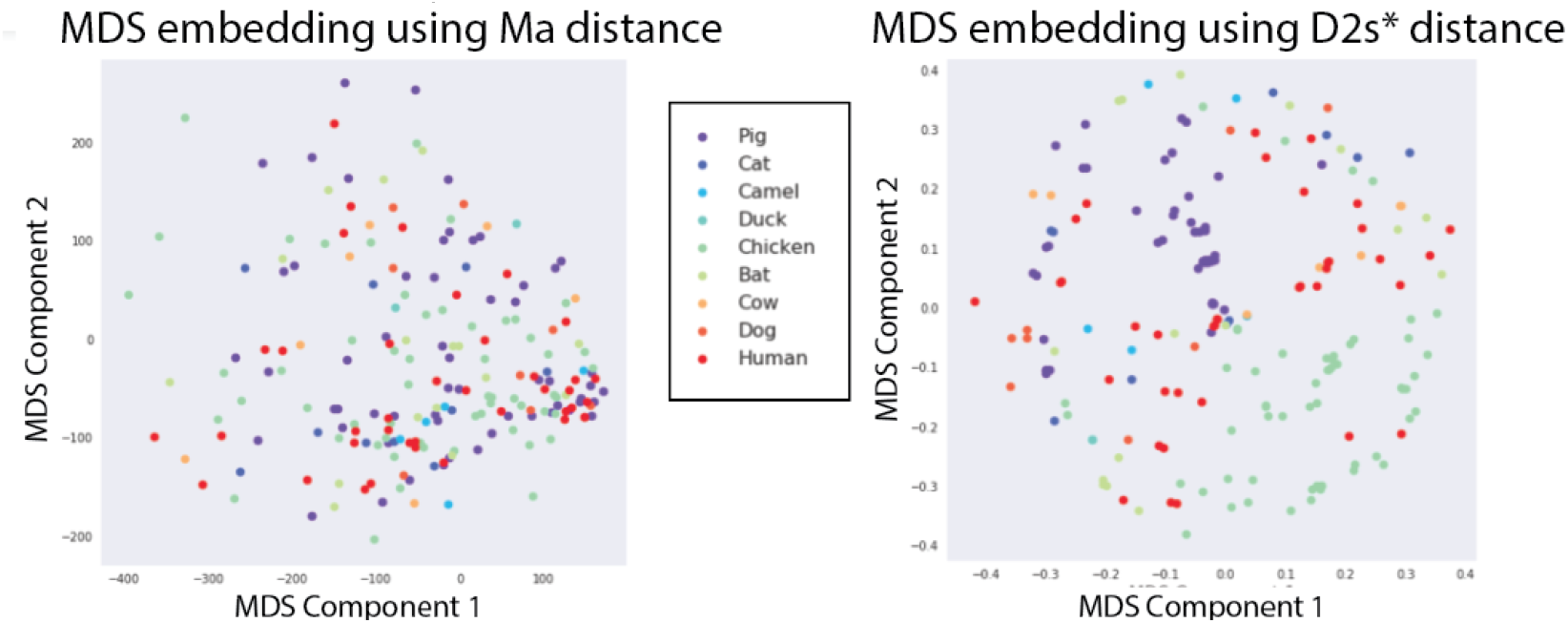
(right): MDS plots for 200 viruses from the Coronaviridae family. Distance computed from the D2*s (right) and Manhattan method (left) on 6-mers profiles.

The Ma metric performed worse with an accuracy of 0.25, far worse than both ViRNN or even KNN with D2S*.

In order to interpret this result, MA and D2*s were visualized by plotting the results of the distance/dissimilarity matrix using multi-dimensional scaling (MDS). As seen on Figure 7 (right), hosts seem to cluster relatively better using the **D2S\*** compared to **Ma**.

As a second model for comparison we implemented a SVM classification task as explained above. In order to compare the result of this SVM with ViRNN the SVM performed classification tasks as binary classification on each class using the OneVersusRest classification method when training the model. As seen in Fig S5, relatively few PCA dimensions are capable of capturing most of the variance in the dataset. We performed optimization of PCA dimension when training the SVM to obtain the best hyperparameter giving the highest accuracy. As a result, the best SVM classification after PCA reduction gave an **accuracy of 84±1**% for a PCA down to 40 components on 6-mers (using 20% of the dataset as testing during 5-cross validation), a **prediction score of 0.91** and an **AUC of 0.96**. The overall accuracy of the SVM is much higher than the KNN, even though the model trained orders of magnitude faster than KNN. Nonetheless, compared to ViRNN the SVM has lower performance: accuracy (84 vs 90%), precision (91 vs 97%) and AUC (0.97 vs 0.96) were all lower compared to ViRNN. When looking specifically at how well the SVM performs for each host, it tends to have better prediction (accuracy) for hosts with large amount of data (Chicken with a accuracy of 98%) while host with smaller amount of data performs the worse (Yak at 0% accuracy, meaning that out of the 26 true Yak hosts in the testing set, none of them were correctly labeled) indicating the SVM was not capable of capturing rare elements. This shows the limitation of the SVM method and the strength of our ViRNN.

### 3.5. Interpretation of RNN

Saliency maps obtained from N-baseline (See Figure S9) had greater range of saliency scores compared to the average nucleotide embedding. Furthermore, background noise in the average embedding was higher compared to the N-baseline. As a result, we preferably interpret the results from the N-baseline that we consider more informative for model prediction.

We generated 3 saliency maps for 3 different viruses that caused 3 seperate epidemics (Covid19, MERS and SARS) by averaging maps from multiple published sequences of the same virus. As shown on Fig. 10, the saliency map reveals peaks in the RNA sequence that are thought to be determinant for the model to make a given prediction (human in this case). Peaks can be ranked based on the saliency score and then mapped onto the genomic sequence to probe into the biological function of a given detected region by this method. The highest peak for SARS-CoV-2 was located in the region coding for the 3C-like proteinase,, a protein implicated in the transcription and replication of the virus ^11^. The second highest peak is located in the coding region of the Spike protein of the virus, a critical protein that enables fusion of the viral membrane with its host. ^12^ Saliency maps on MERS and SARS (Fig. S10,11) both featured their highest peak in the region coding for the non structural protein 3 (nsp3), an essential protein in the transcription machinery of the two viruses. ^13^

**Figure 8:**
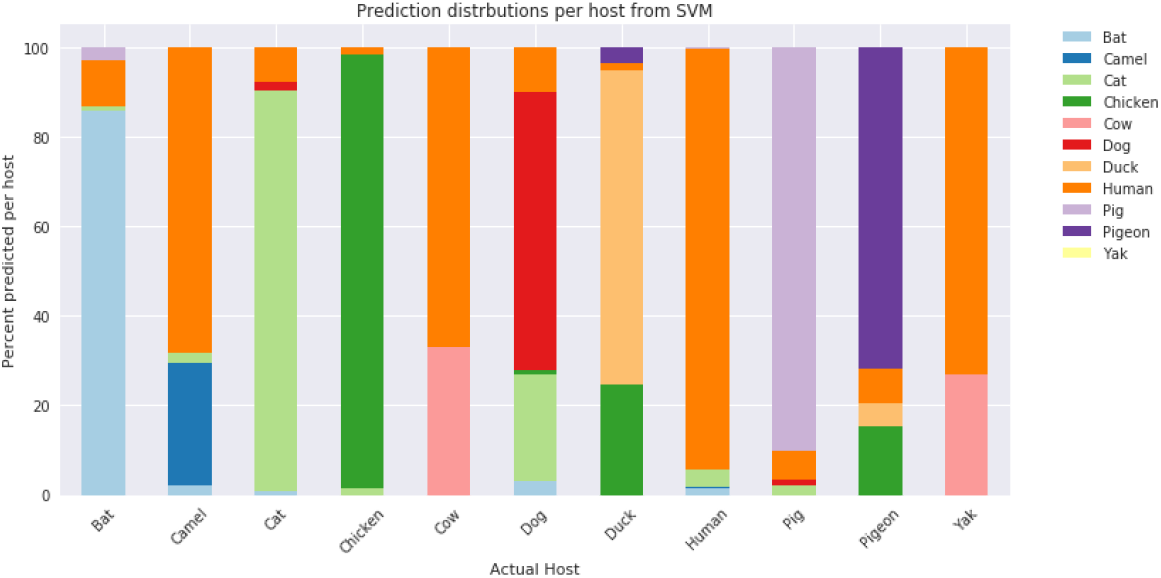
Model prediction for each host using the SVM classification method on the validation set.

**Figure 9:**
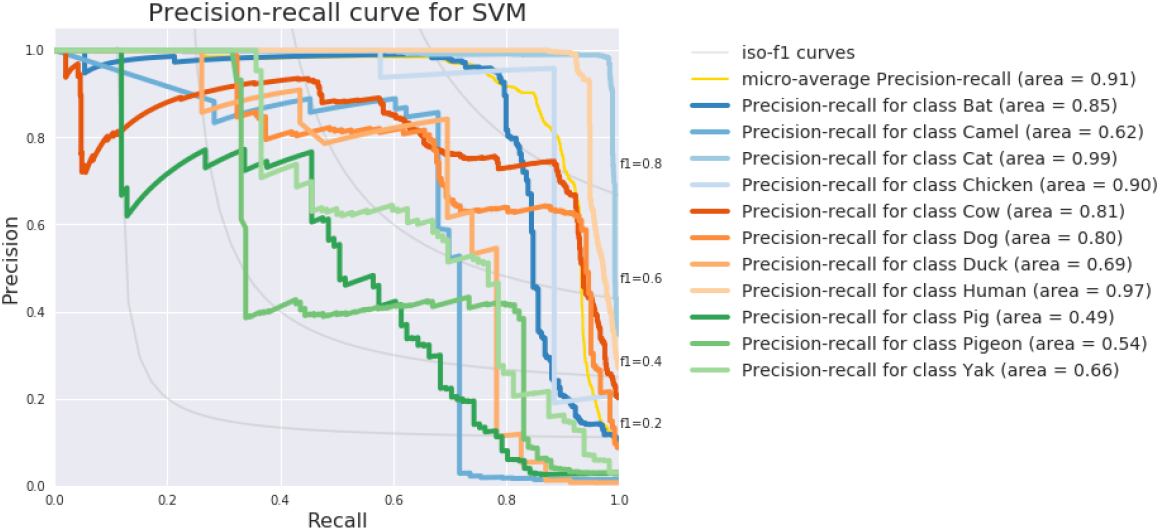
Precision-Recall Curve for SVM classification method on validation set. Each curve corresponds to binary classification.

**Figure 10:**
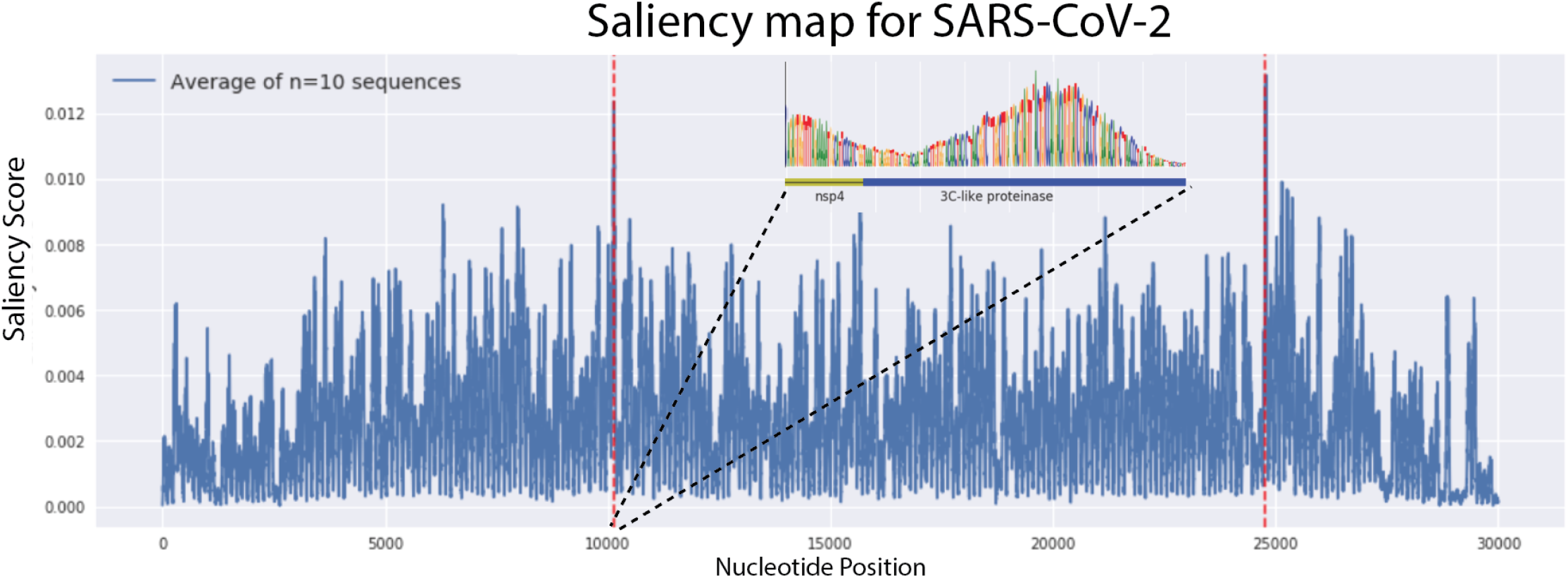
Saliency map of the SARS-CoV-2 genome from the ViRNN. Map averaged over 10 sequences. Saliency score was computed using integrated gradient method from a N-baseline (‘any nucleotide’). Zoom into the [10020-10090] nucleotide window features nucleotide logos of important sequences with alignments with the translated protein

## 4. Discussion

### 4.1. Model Training

In this project we have trained a combined convolutional and recurrent neural network to predict the host of a virus from genomic sequence. Although we derived our model architecture from recently published work in this area, we tested a couple of different architectures including LSTM and GRU recurrent layers, as well as varying the input sequence length to test the best parameters for our task. LSTM and GRU layers are commonly used in networks when analyzing sequence grammar, particularly in written text or genomic sequences. We initially implemented LSTM layers as previously published but wanted to speed up training so we could test multiple models. GRU layers have recently been used in place of LSTMs because they perform similarly but have a reduced number of parameters because they lack a memory unit ^14^. We expected this to decrease our training time, which it did marginally (not quantified), but instead, we found the GRU architecture performed much better on this task in terms of the validation loss after 5 epochs. In fact, after 5 epochs our model had a categorical accuracy of almost 78%, and only required an additional 2 epochs of training to reach the final model described in this paper. Other reports have demonstrated increased performance of GRUs compared to LSTM on genomic sequence data ^15^. Although this may be an artifact of faster training, in our hands we were never able to train an LSTM network to match the performance of the GRU based networks for this task. This may suggest some benefit of using GRUs over LSTMs in the future for extracting features from genomic sequence information.

Our input dataset spanned eleven different hosts, and while neural networks usually perform well on this type of classification task, we were concerned about the class imbalance in our data. We trained the network with class weightings in order to correct for potential biases between the number of sequences per host, and trained with a batch size of 500 to ensure that even the least represented samples would be present in a random batch drawing of the dataset. Our initial models had some issues with overpredicting the most abundant hosts like human and chicken, however, it appears in the final model that the class imbalance didn’t dictate all of the error. For example, Yak coronaviruses were the least represented in our dataset, yet our model was capable of predicting the correct host for 85-90% of these viruses. On the other-hand, the model struggled with dog coronaviruses, which could have to do with some sequence characteristics of the viruses rather than the model itself. Future work will need to be done to elucidate what makes some of these hosts harder to predict than others.

During model training we also came across the interesting observation that the model performed better on full length sequences rather than the overlapping subsequences on which we trained ViRNN. Previous studies haven’t witnessed this phenomenon, although this is a relatively new field. We speculate this could have to do with having more information per sequence with which to make a decision, but further interpretability work could help us understand this observation.

We also tested this model on the *Orthomyxoviridae* family of viruses, of which Influenza is a member. It was not so surprising that the model performed poorly on this dataset as it is out of distribution and biologically, this family is quite different from coronaviruses. We initially desired to test this dataset on the *Arenavirales* family of viruses, which is phylogenetically closer to coronaviruses, but were not able to due to little host overlap between Coronaviruses and this family. Further exploration could certainly be done with viruses that have structure more similar to coronaviruses to see if performance is increased.

### 4.2 KNN vs. SVM vs. RNN

We implemented canonical ML methods to predict hosts from unaligned viral sequences, namely KNN and SVM. Both of these methods rely on representation of a genomic sequence into a kProfile using k-mer representation, therefore limiting the ability to interpret the result because they do not retain spatial information of the words of nucleotides. The distance metric used in KNN is a key determinant of performance. Even though the elaborate dissimilarity metric D2S*, which requires Monte Carlo simulations for k-mer probability distribution ^2,8^, performed dramatically better than common metrics (Ma), the computational cost of this method makes it prohibitive to use it on large datasets and is not adequate for big data. Published methods only report from 700 up to 1000 sequences comparison at a time ^2^. SVM classification on the other hand had much better success compared to KNN but in all of the hyperparameter that we tested (length of k-mers, components for PCA reduction) the accuracy of this method capped at 84% and performed mediocrely on out of distribution hosts (cow and yak). Another hyperparameter that could be tested using SVM classification is the type function used as kernel. For this work we relied on previous published work ^2^ invoking superiority of the Radial Basis Function over other kernels (such as polynomial or sigmoid). Nevertheless, little information can be extracted after the SVM model has been trained due to the fact that the sequences were converted into kProfiles and thereafter reduced into fewer components by PCA reduction. SVM also failed to classify rare sequences in our dataset. As a result the SVM method fails in two ways compared to our ViRNN: inferiority in terms of accuracy and lack of ability to be interpreted in a biological context.

### 4.3. Model Interpretation

Our initial tries to generate saliency maps using integrated gradients were unsuccessful, as it appears the highest gradients are simply found at the end of the sequence. This likely is a vanishing gradient artifact generated from the bidirectional GRU modules in the model. Further work could be done to optimize the ViRNN architecture to prevent this issue.

We utilized a modified protocol to generate saliency maps by stitching together maps from smaller, mostly overlapping, subsequences of data. This allowed us to generate saliency maps for three important coronaviruses, SARS, SARS-CoV-2, and MERS.

When looking at the regions with the largest saliency score, we see that they lie in regions including the 3C-like proteinase and the spike protein. The 3C-like proteinase is responsible for cleaving the polyproteins generated from the translation of the viral genome into the constituent proteins that go on to promote viral replication^16^. This is key for the survival of the virus inside the host cell. It is possible that certain host factors affect the ability of this protease to function, thus conferring a selective pressure between different hosts to mutate this protein. Diversity in this protein amongst coronaviruses is poorly characterized, future work will need to look into this area as a potential way of determining the host of a virus.

We also found a region in the S2 domain of the spike protein to be among the most important in terms of saliency score. This protein is key for entry of the virus, and has two different cleavage sites, one between the S1/S2 boundary, and another inside the S2 domain^17^. These are key for entry into the cell. The area of interest identified did not lie within either of these two cleavage sites, and thus could represent a new site that is important for host tropism. It is difficult to draw concrete conclusions from this though, as the biological relevance is unknown for this particular site in the protein.

Another interpretation method known as SHAP (SHapley value Additive exPlanations) relying on game theory was successfully implemented for RNN^18^ recently and could also provide valuable information about the underlying mechanisms of ViRNN for host prediction. The field of deep learning interpretation^19^ is still fairly new and we expect that valuable information will be extracted from such methods when implemented on newly established models like ViRNN.

### 4.4. Future Directions

Moving forward there are a number of angles to explore. Although this model works well for coronaviruses, it doesn’t appear to have predictive power for other virus families. Further work might seek to identify an ideal training set that would expand the predictive capabilities of the model while still allowing accurate host prediction. For example, there might be an optimal set of testing sequences across multiple hosts and viral species. Because the model outputs probabilities for each host, it may be possible to use these model outputs as a predictor for zoonotic transfer between species. We were not able to explore this in our project given the time and dataset available, but a curated dataset of sequences with known zoonotic transfer events may allow this idea to be further explored.

In our model we utilized genomic sequence information on the raw sequence level, however, further work could also be done to incorporate protein level information into this model. Perhaps addition of protein level information could lead to more interpretable decisions if the model is able to delineate particular proteins and their importance to the tropism of the virus.

Overall, we present a deep learning method for accurately identifying viral hosts from coronavirus genome sequences. This has applications in virology and could be used in the future in epidemiology to understand zoonoses.

## Acknowledgements

We would like to thank Sachit Saksena and Corban Swain for their helpful discussions and suggestions.

## 6. Supplementary data

**Figure S1:**
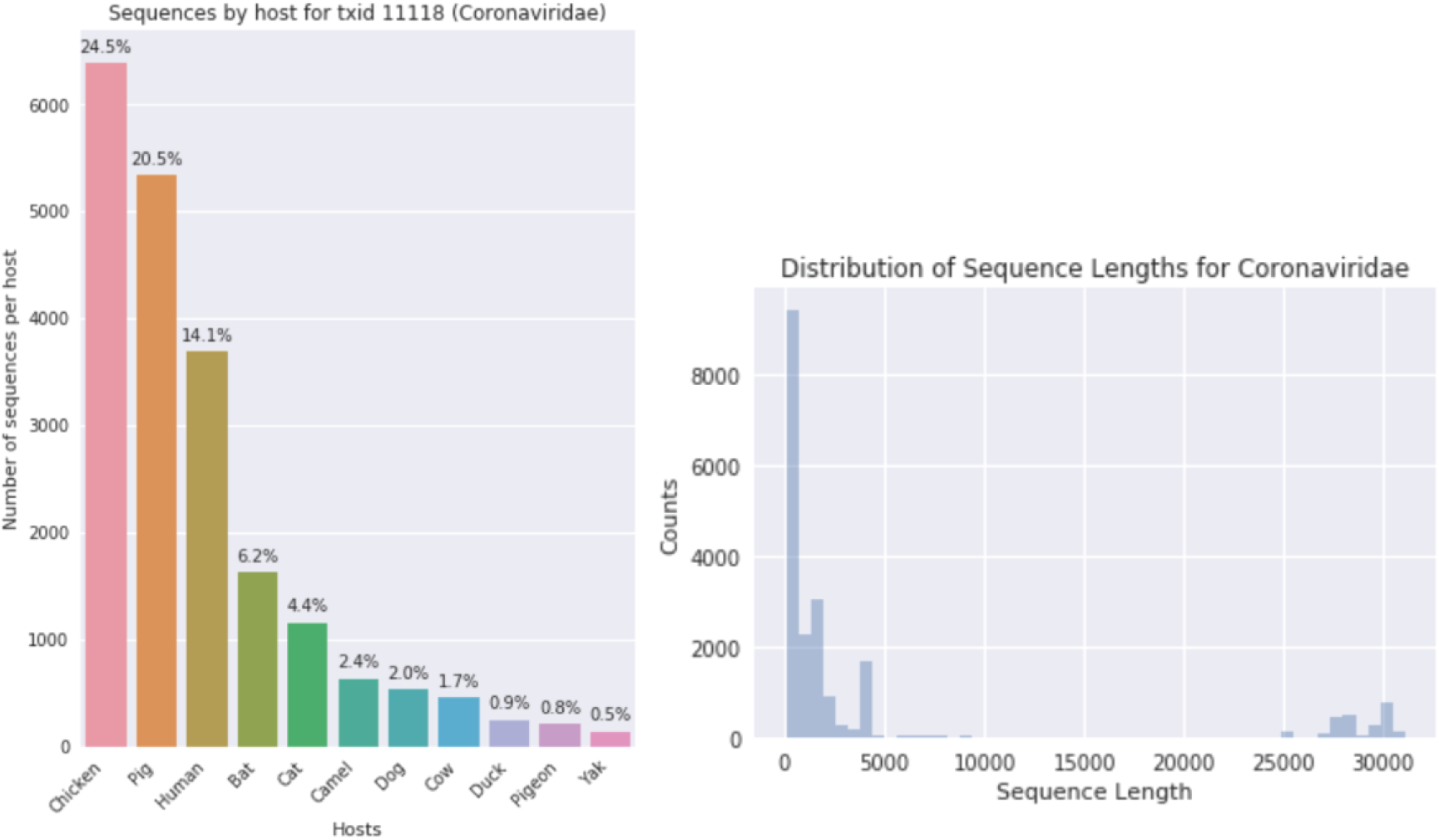
Host and sequence length distribution of *Coronaviridae* family. (LEFT) Number of sequences per host, with percentage of total sequences listed per host. (RIGHT) Distribution of sequence length for all sequences.

**Figure S2:**
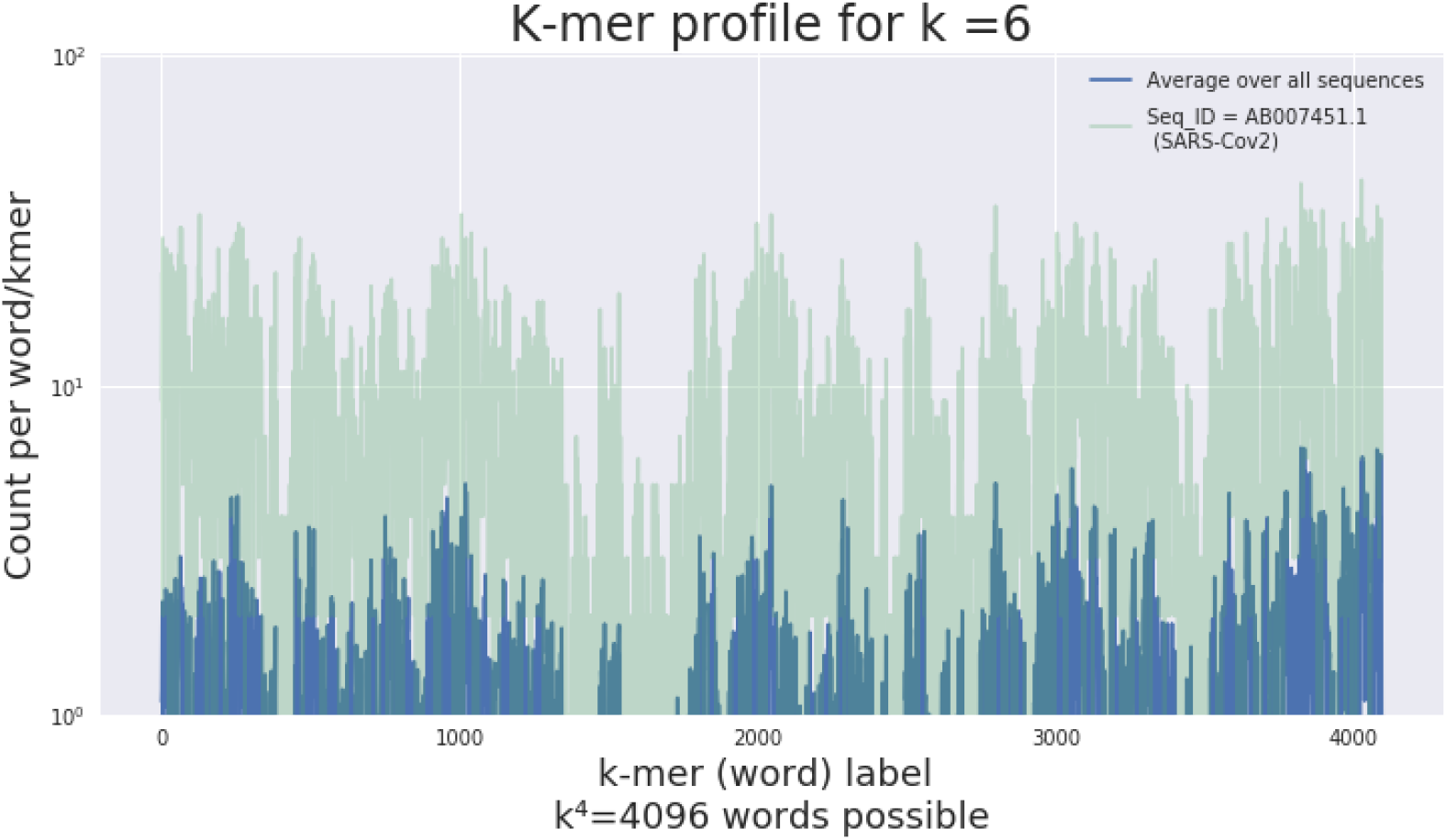
kProfile (k=6) for the *Coronaviridae* family. Average over all sequences (blue) and example of one sequence for SARS-Cov-2 (green)

**Figure S3:**
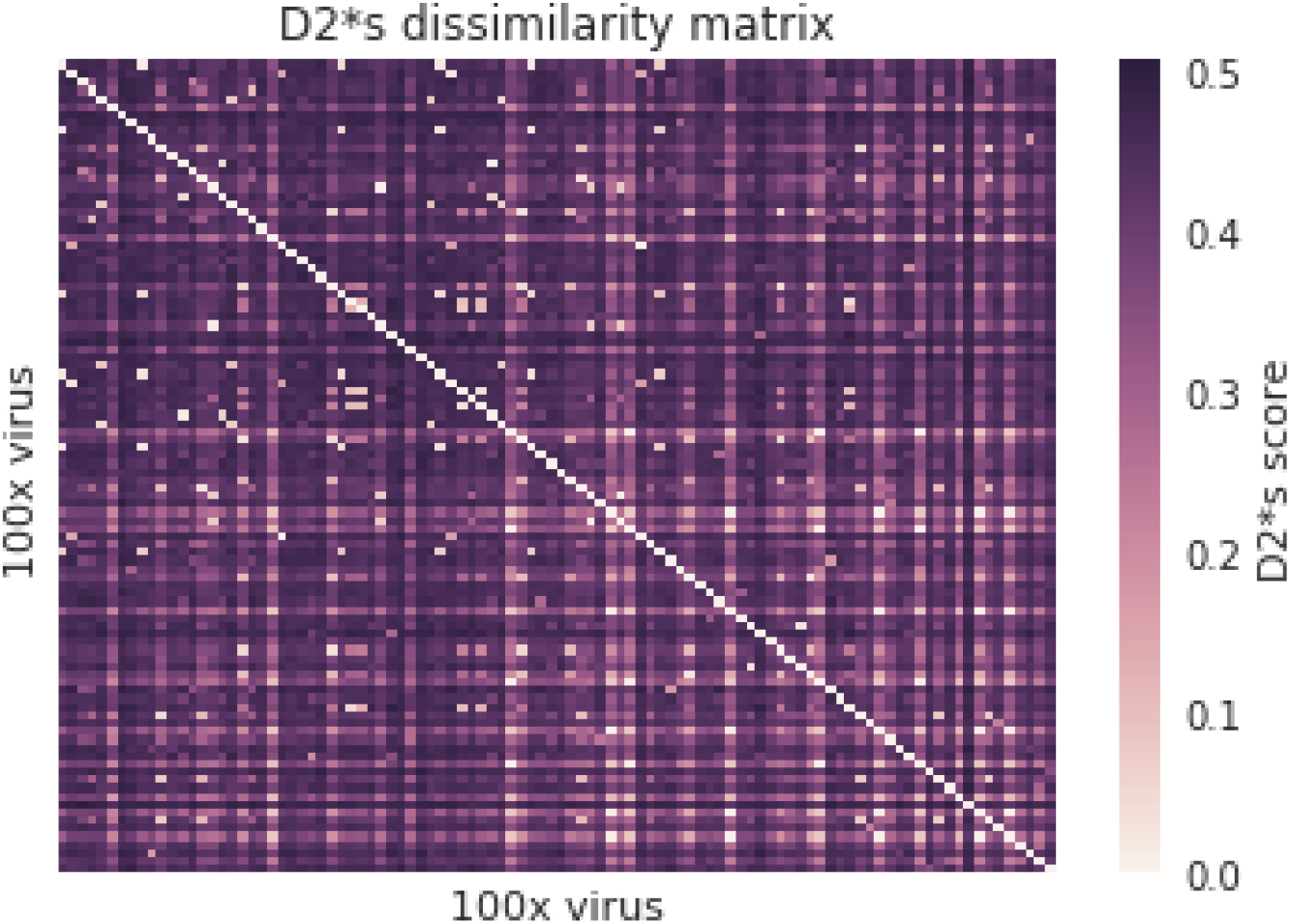
Dissimilarity D2*s matrix on 100x samples from the *Coronaviridae* family. Each column represents one virus, each row represents its D2*s score against the other viruses.

**Table S4:** Computational time for dissimilarity matrix D2S* for KNN classification on 6-mer Profiles. Configuration used was 16 parallelized CPUs.

| Matrix size | Computational time |
| --- | --- |
| 10 x 10 | 2 min |
| 100 x 100 | 41 min |
| 200 x 200 | 112 min |
| 500 x 500 | 382 min |
| 26 066 x 26 066 | Not computed - Expected time > 15 days |

**Figure S5:**
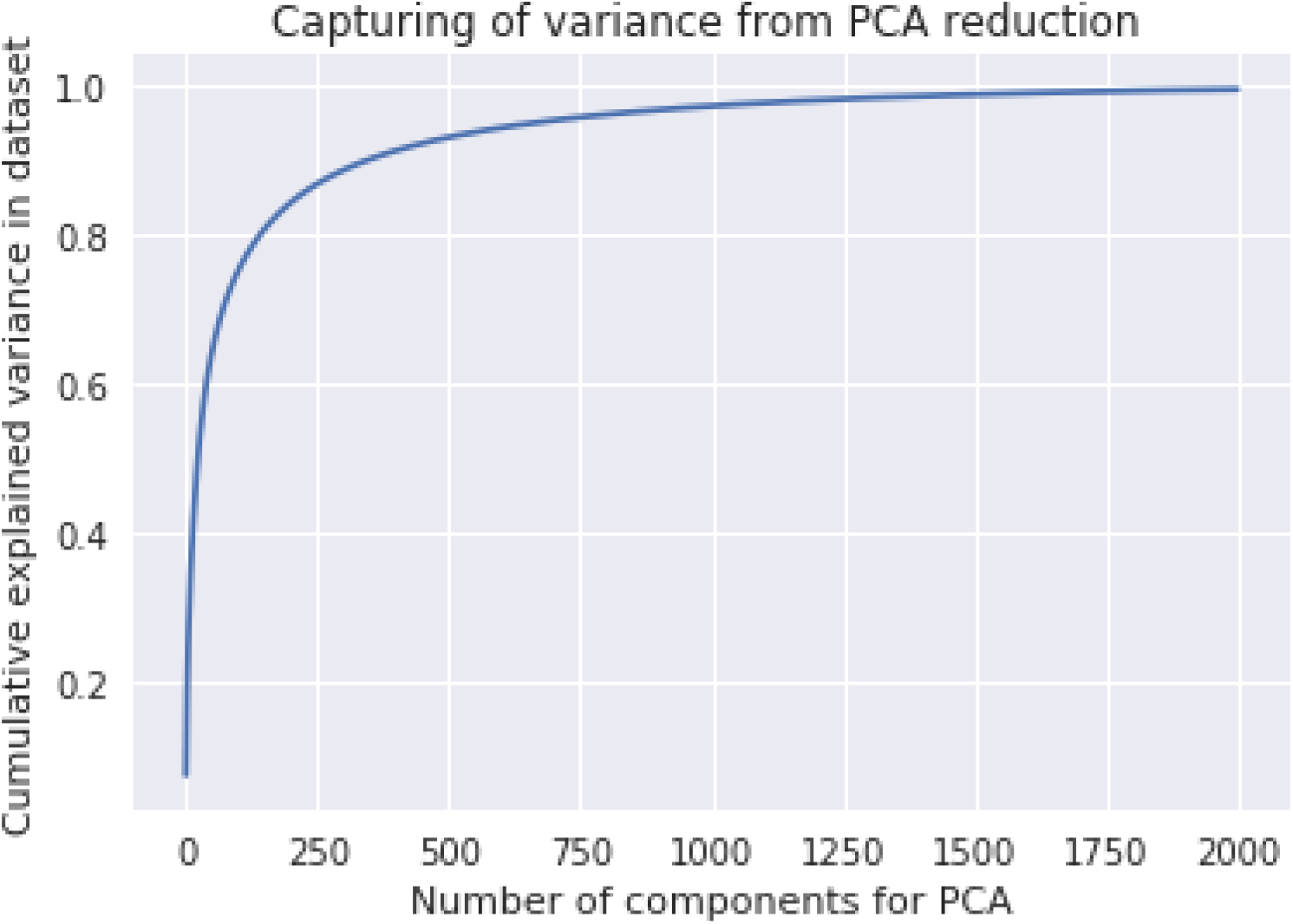
Cumulative variance in *Coronaviridae* dataset from 6-kProfiles from PCA reduction. More than 80% of the cumulative variance is conserved with only 140 components

**Figure S6:**
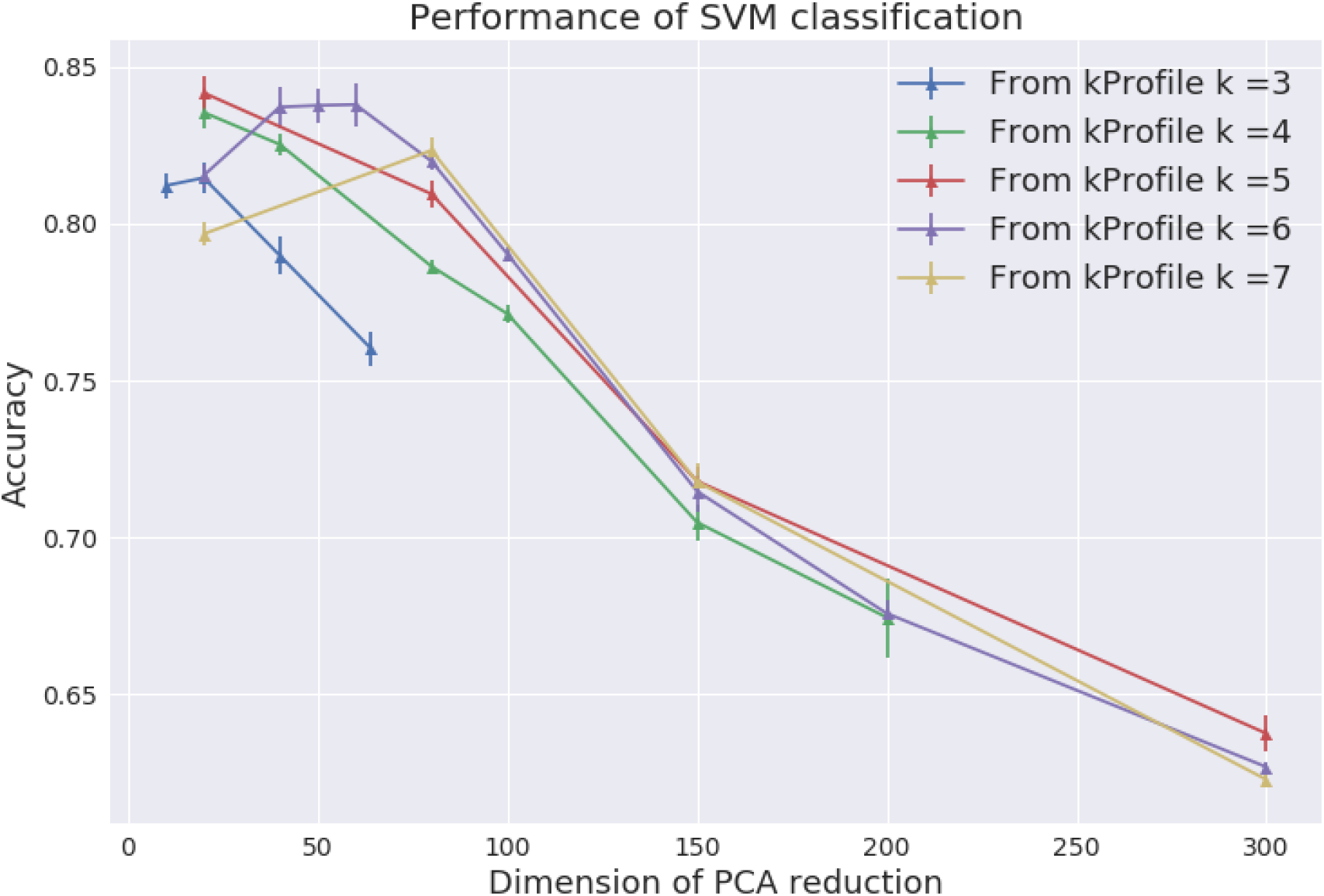
Accuracy of the SVM method for host classification on the *Coronaviridae* family. Accuracy computed for multiple PCA dimension reduction and kProfiles. Accuracy on the testing set using a 5-cross validation method on the entire dataset. For 3Profile, number of 3-mers = 64 (blue).

**Figure S7:**
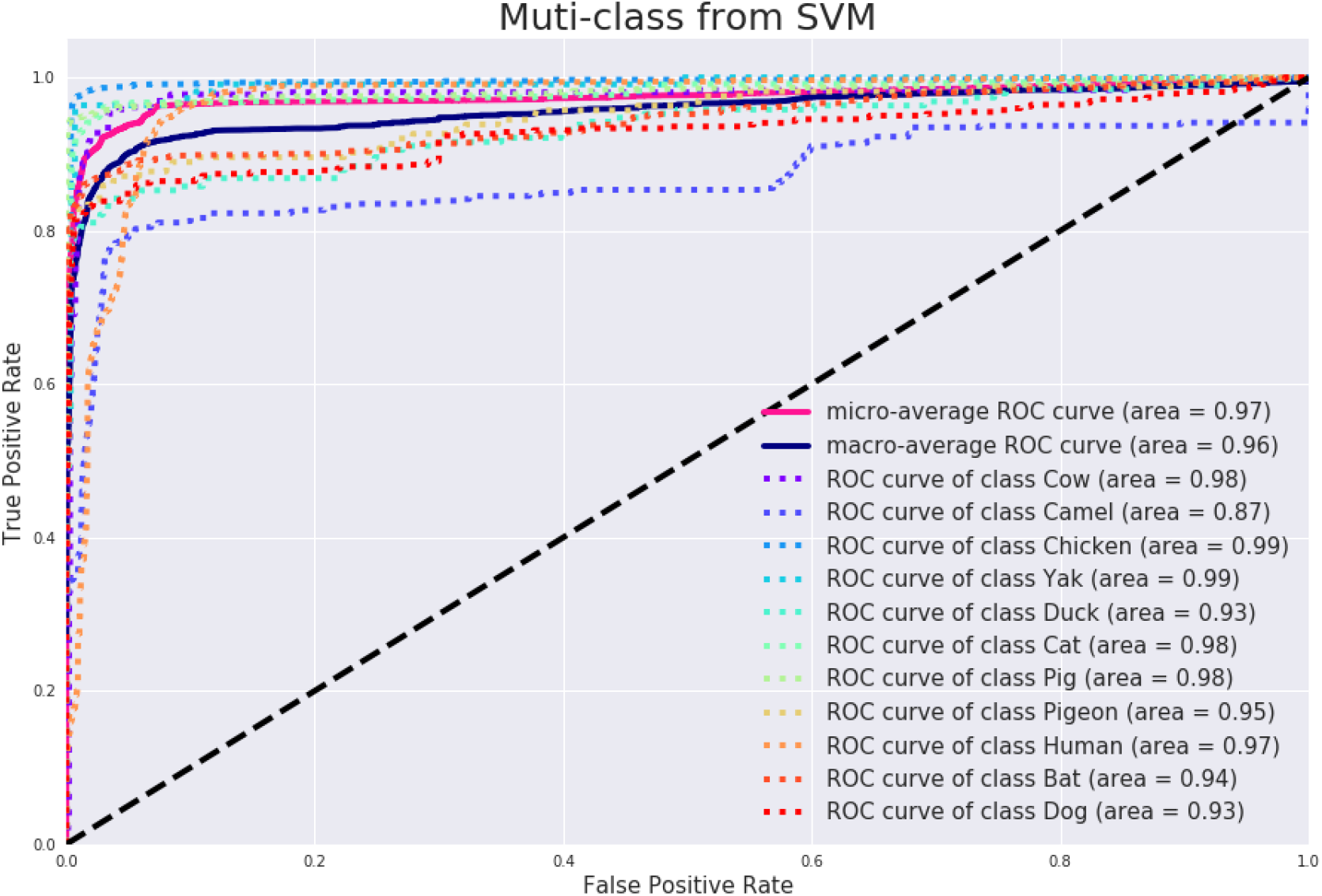
ROC curve for the SVM classification method using 6-mers Profile. Curve corresponding for binary classification and for micro and macro classification.

**Figure S8:**
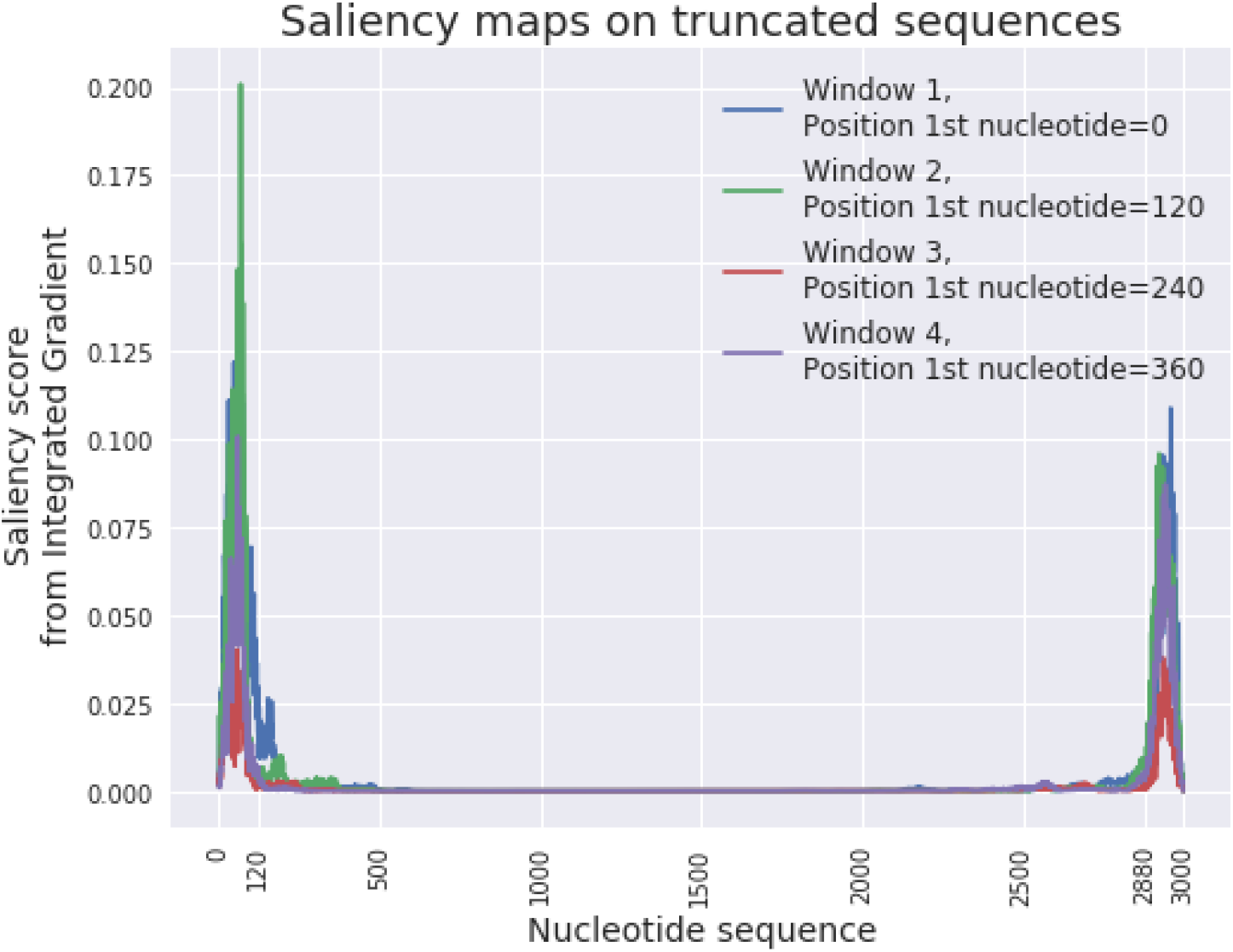
Saliency maps on truncated sequences from sliding windows for a given viral sequence.

**Figure S9:**
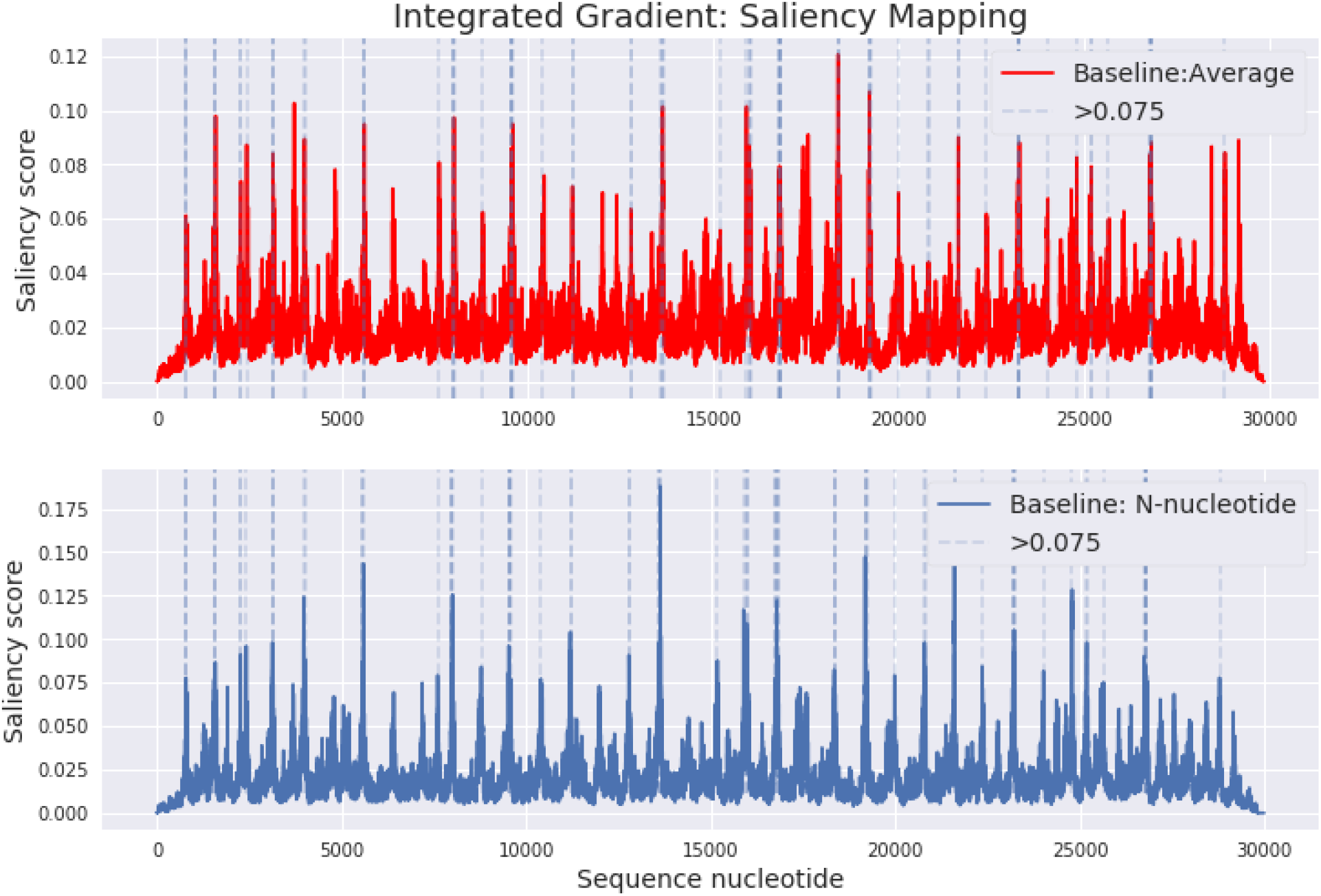
Saliency map from integrated gradient method on an individual genomic sequence from the SARS-CoV-2 virus. <u>Top:</u> map computed with an Average nucleotide embedding baseline, <u>Bottom:</u> from N-nucleotide baseline. Dashed blue lines indicate peaks above >0.075 threshold for the N-baseline. Most peaks from the N-nucleotide baseline are conserved on the Average baseline. Peak values of saliency score are higher in the N-baseline compared to the peaks from the Average baseline.

**Figure S10:**
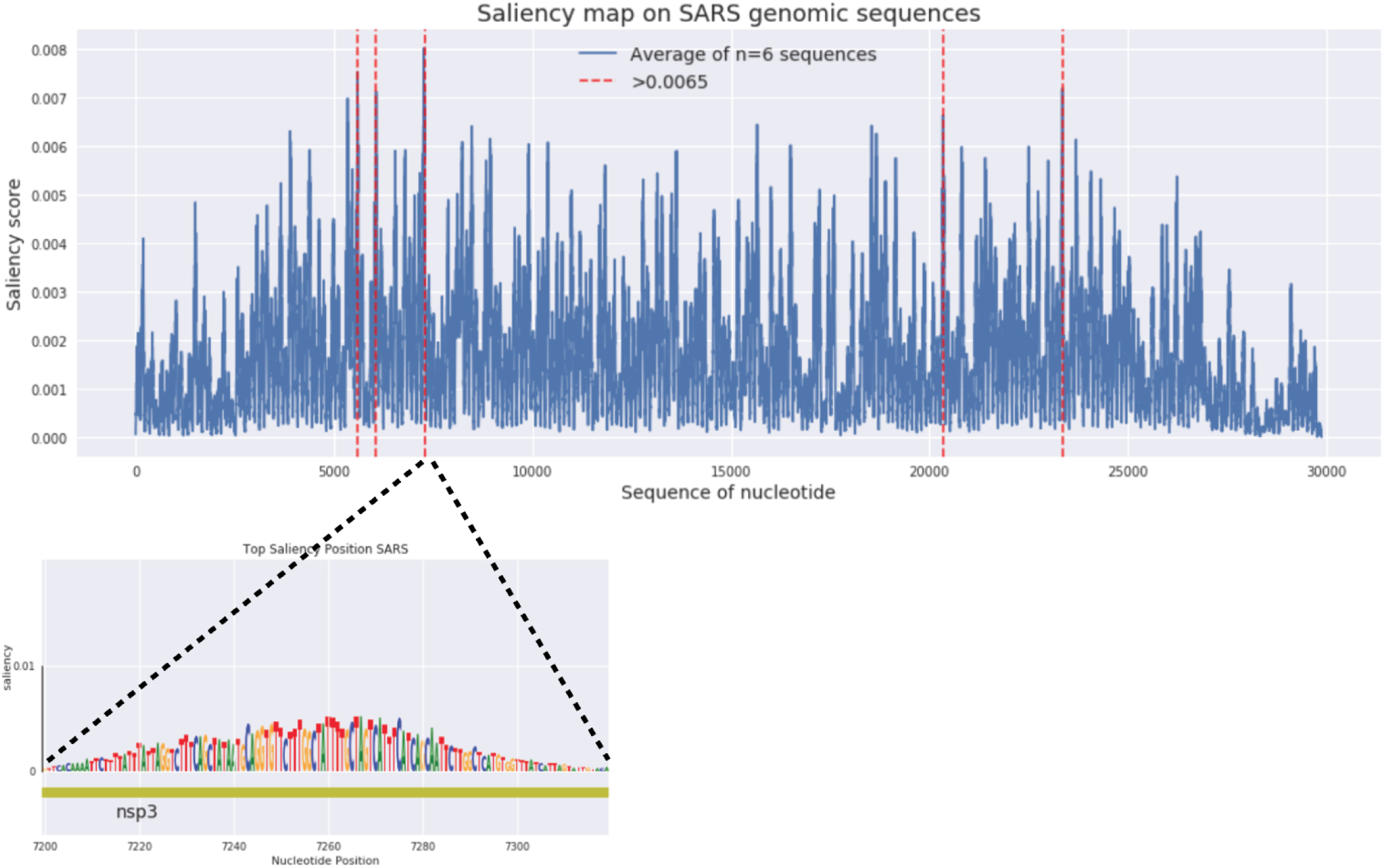
Saliency map of the SARS genome from the ViRNN. Map averaged over 6 sequences. Saliency score was computed using the integrated gradient method from a N-baseline (‘any nucleotide’). Zoom into the [7200-7380] nucleotide window features nucleotide logos of important sequences with alignments with the translated protein

**Figure S11:**
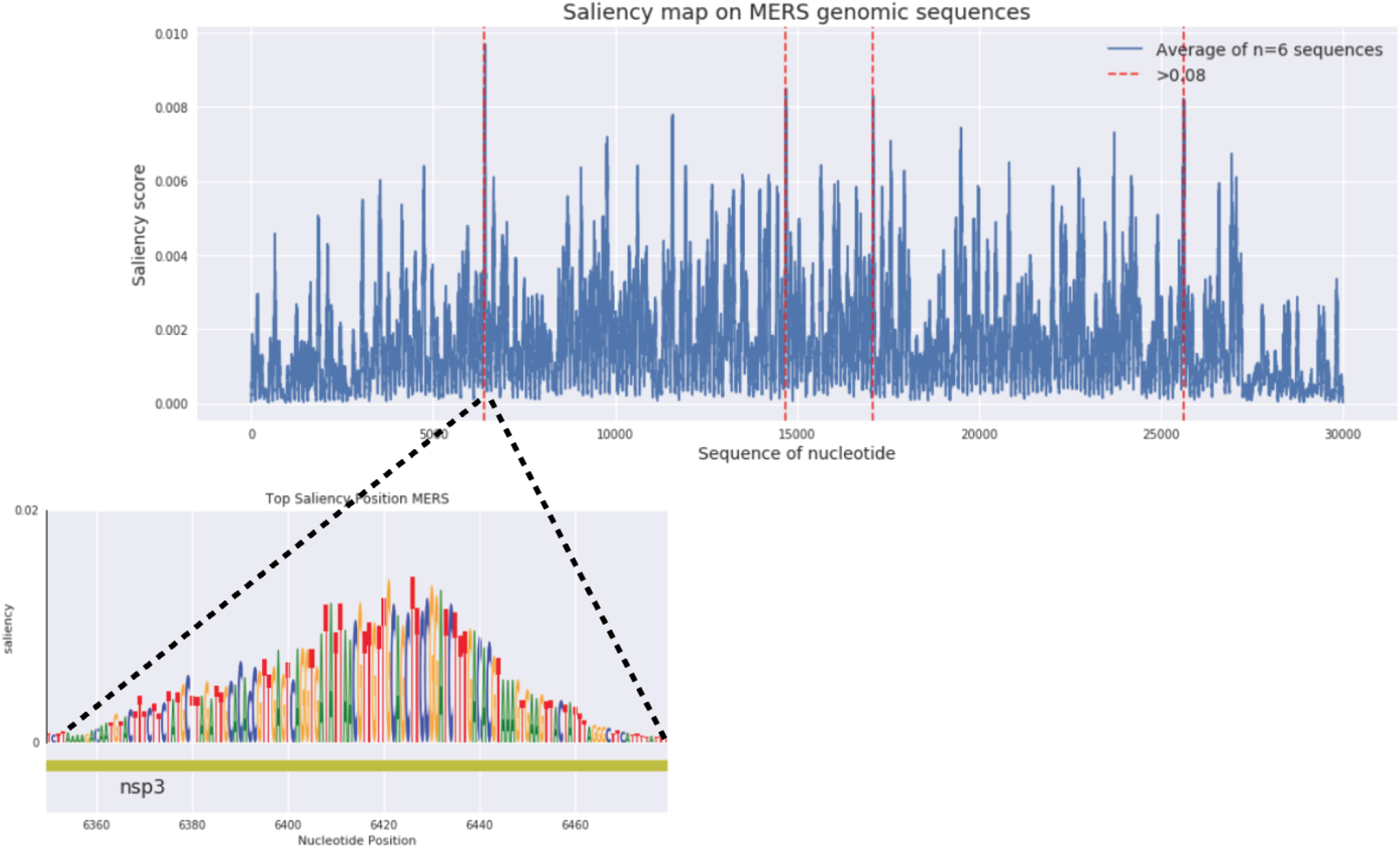
Saliency map of the MERS genome from the ViRNN. Map averaged over 6 sequences. Saliency score was computed using the integrated gradient method from a N-baseline (‘any nucleotide’). Zoom into the [6355-6476] nucleotide window features nucleotide logos of important sequences with alignments with the translated protein

**Figure S12:**
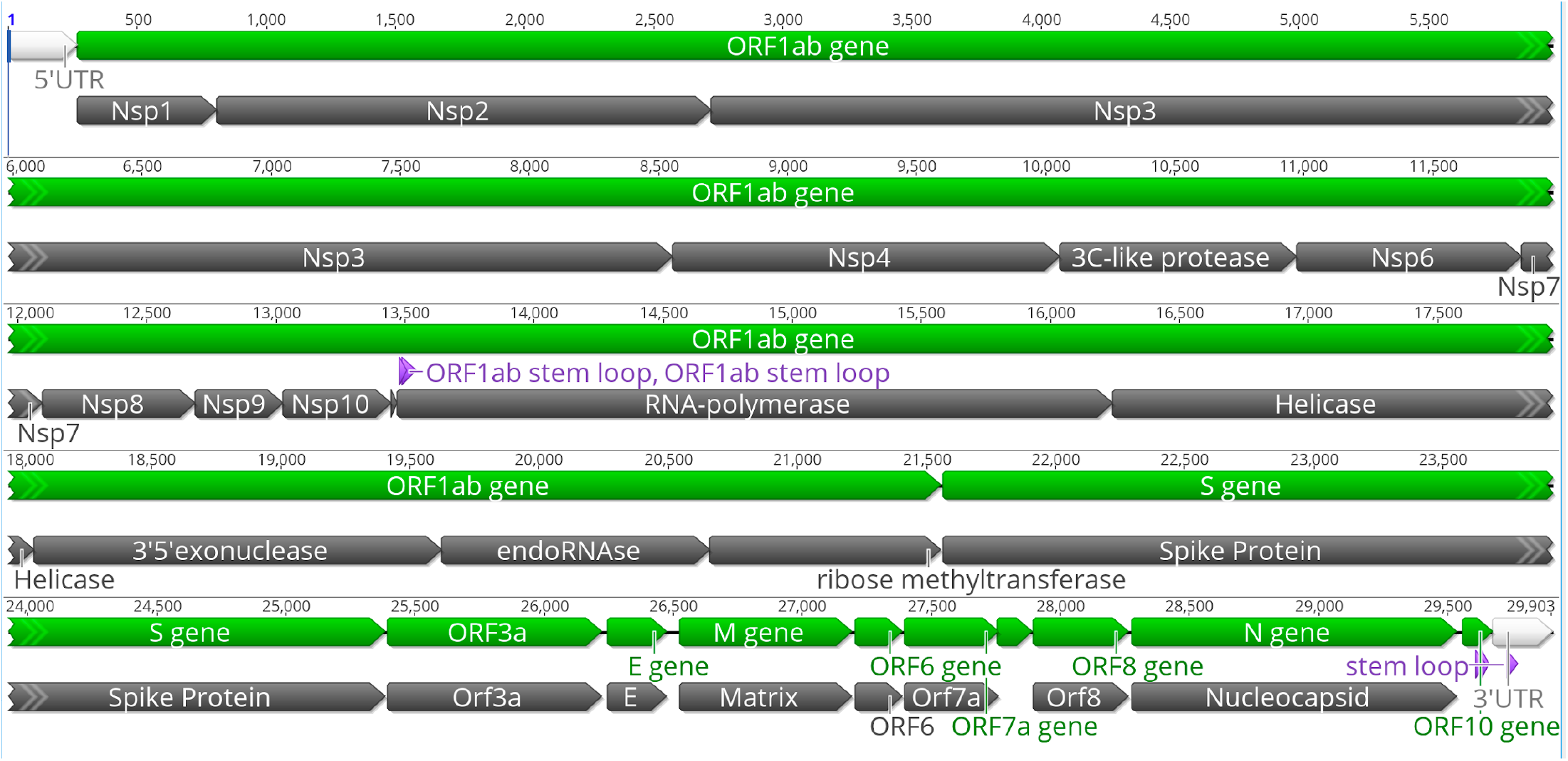
RefSeq genome of SARS-CoV-2 (NC_045512). Line representing nucleotide sequence, green indicating translated genes and black translated polypeptide or protein from the virus.

## Notes

### Competing Interest Statement

The authors have declared no competing interest.

